# The hydrophobin-like OmSSP1 may be an effector in the ericoid mycorrhizal symbiosis

**DOI:** 10.1101/227959

**Authors:** Salvatore Casarrubia, Stefania Daghino, Annegret Kohler, Emmanuelle Morin, Hassine-Radhouane Khouja, Yohann Daguerre, Claire Veneault-Fourrey, Francis M. Martin, Silvia Perotto, Elena Martino

**Author notes:** **Correspondence** Elena Martino.

## Abstract

Mutualistic and pathogenic plant-colonizing fungi use effector molecules to manipulate the host cell metabolism to allow plant tissue invasion. Some small secreted proteins (SSPs) have been identified as fungal effectors in both ectomycorrhizal and arbuscular mycorrhizal fungi, but it is currently unknown whether SSPs also play a role as effectors in other mycorrhizal associations. Ericoid mycorrhiza is a specific endomycorrhizal type that involves symbiotic fungi mostly belonging to the Leotiomycetes (Ascomycetes) and plants in the family Ericaceae. Genomic and RNASeq data from the ericoid mycorrhizal fungus *Oidiodendron maius* led to the identification of several symbiosis-upregulated genes encoding putative SSPs. OmSSP1, the most highly symbiosis up-regulated SSP, was found to share some features with fungal hydrophobins, even though it lacks the Pfam hydrophobin domain. Sequence alignment with other hydrophobins and hydrophobin-like fungal proteins placed OmSSP1 within Class I hydrophobins. However, the predicted features of OmSSP1 may suggest a distinct type of hydrophobin-like proteins. The presence of a predicted signal peptide and a yeast-based signal sequence trap assay demonstrate that OmSSP1 is secreted during symbiosis. OmSSP1 null-mutants showed a reduced capacity to form ericoid mycorrhiza with *Vaccinium myrtillus* roots, suggesting a role as effectors in the ericoid mycorrhizal interaction.

## Introduction

Fungi secrete a wide range of enzymatic and non-enzymatic proteins that function in the break-down of complex organic molecules but also in the interaction with microbial competitors or with animal and plant partners (Scherlach et al., 2013; Stergiopoulos & de Wit, 2009; Talbot et al., 2013; Tian et al., 2009). Fungi can establish different types of interactions with plants, ranging from mutualistic to antagonistic. Whatever their lifestyle, plant-colonizing fungi are recognized by the plant immune system through invariant molecular patterns known as microbe- or pathogen-associated molecular patterns (MAMPS or PAMPs) (Jones & Dangl, 2006). To successfully colonize plant tissues, fungi must prevent the PAMP-triggered immunity (PTI) reaction (Lo Presti et al., 2015). For this purpose, fungi secrete effector molecules that may play different functions depending on the fungal lifestyles. For example, they can be toxic compounds that kill the host plant (in necrotrophs), or secreted proteins that shield the fungus and suppress the host immune response, or proteins that manipulate the host cell metabolism to allow plant tissues invasion and nutrient uptake (de Jonge et al., 2011; Giraldo et al., 2013; Selin et al. 2016). Many small secreted proteins (SSPs) have been reported to function as effectors (Lo Presti et al., 2015).

Effectors were initially considered as virulence factors secreted exclusively by pathogens (Stergiopoulos & de Wit, 2009; van Esse et al., 2008). However, it has become apparent that effectors can manipulate the plant immune system also in mutualistic associations (Kim et al., 2016). Mutualistic fungi establish intimate contacts with plants by forming specialised fungal structures involved in nutrient exchange with the host (Martin et al., 2017). Effector-like SSPs have been functionally characterized as effector-like molecules both in arbuscular (AM) and ectomycorrhizal (ECM) fungi, as well as in some endophytic fungi (Plett & Martin, 2015). For example, the ECM fungus *Laccaria bicolor* requires MiSSP7 (Mycorrhizal induced Small Secreted Protein 7) to establish symbiosis. MiSSP7 suppresses the plant defence reactions by interacting with the jasmonate co-receptor JAZ6 (Plett et al., 2011; 2014). Similarly, the AM fungus *Rhizophagus irregularis* secretes SP7, an effector protein that counteracts the plant immune program by interacting with the pathogenesis-related transcription factor ERF19, leading to increased mycorrhization (Kloppholz et al., 2011). Tsuzuki et al. (2016) also showed that host-induced gene silencing of SlS1, a putative secreted *R. irregularis* SSP expressed in symbiosis, resulted in suppression of colonization and formation of stunted arbuscules. In a similar manner, the candidate effector protein PIIN_08944, secreted by the fungal endophyte *Piriformospora indica* during colonization of both *Arabidopsis thaliana* and barley plants, was demonstrated to play a crucial role, in reducing the expression of PTI genes and of the salicylic acid defense pathway (Akum et al., 2015).

Protein effectors are normally secreted following the endoplasmic reticulum-Golgi apparatus system, and bioinformatic identification of effector candidates can thus be based on the presence of the N-terminal signal peptide (Lo Presti et al., 2015), even though alternative secretion pathways have been reported for *Magnaporthe oryzae* and *Phytopthora infestans* (Giraldo et al., 2013; Wang et al. 2017). General SSPs features are: a) the presence of a signal peptide and the absence of transmembrane domains or GPI-anchor sites; b) a small size, with a mature length smaller than 300 amino acids; c) a richness in cysteine residues and, sometimes, d) the presence of conserved motifs (Hacquard et al., 2012; Lo Presti et al., 2015; Martin et al., 2008; Stergiopoulos & de Wit, 2009; Zuccaro et al., 2014). Bioinformatic analyses of about fifty fungal genomes has highlighted that, when compared with saprotrophic and pathogenic fungi, the ECM fungal secretome is enriched in SSPs and contains species-specific SSPs likely dedicated to the molecular cross-talk between fungal and plant partners (Pellegrin et al., 2015). In line with this finding, a comparative *in silico* analysis of the AM fungi *Rhizophagus clarus*, *R. irregularis* and *Gigaspora rosea* highlighted the presence of shared SSPs (Sędzielewska Toro & Brachmann, 2016; Kamel et al., 2017), supporting a general conserved role of SSPs in AM. These data suggest that effector SSPs may represent an important fungal “toolkit” that enables the establishment/maintenance of host plant colonization in mycorrhiza (Plett & Martin, 2015, Martin et al., 2016). However, there is increasing awareness that SSPs in mycorrhizal fungi are likely involved in additional functions unrelated to symbiosis. For example, several SSPs are secreted by the ECM fungi *L. bicolor* and *Hebeloma cylindrosporum* during the free-living phase (Doré et al., 2015; Vincent et al., 2012). Moreover, large-scale transcriptomic and genomic analyses including fungi with different lifestyles revealed a wide array of SSPs in most saprotrophic fungi (Pellegrin et al., 2015; Valette et al., 2017), suggesting a possible role for SSPs in competition and rhizospheric communication (Rovenich et al., 2014).

Whereas effector SSPs have been characterized in AM and ECM fungi, there is currently no information on the occurrence of SSPs in ericoid mycorrhizal (ERM) fungi and on their potential role in symbiosis. ERM fungi are soil-borne fungi mostly belonging to Leotiomycetes (Ascomycetes). They form a peculiar endomycorrhizal type by colonizing the root epidermal cells of plants within the family Ericaceae and promote growth of their host plant in stressful habitats (Perotto et al., 2012). A common ERM fungal species is *Oidiodendron maius* (Dalpé, 1986) and *O. maius* strain Zn, an isolate from a metal polluted soil whose genome and transcriptome have been recently sequenced (Kohler et al., 2015), has become a model system to investigate metal stress tolerance in these fungi (Daghino et al., 2016; Perotto et al., 2012; Ruytinx et al., 2016). Genomic data have revealed a large set of carbohydrate-active enzymes (CAZymes) in *O. maius,* with many plant cell wall degrading enzymes being expressed during symbiosis (Kohler et al., 2015). As these enzymes could potentially elicit defense reactions through oligosaccharide release, symbiosis development likely requires a tight control of the plant defense reactions and, based on our current knowledge of arbuscular and ectomycorrhizal interactions, effectors to control plant immunity. Aim of this work was to identify, through the analysis of *O. maius* genomic and transcriptomic data, fungal SSPs potentially involved in the molecular dialogue governing the ERM symbiosis.

## Materials and Methods

### Fungal strains and growth conditions

*Oidiodendron maius* strain Zn (hereafter *O. maius*) was isolated from the roots of *V. myrtillus* growing in the Niepolomice Forest (Poland), and first described by Martino et al. (2000). This *O. maius* strain is deposited at the Mycotheca Universitatis Taurinensis collection (MUT1381; University of Turin, Italy) and at the American Type Culture Collection (ATCC MYA-4765; Manassas, VA, US), and was maintained on Czapek-Dox solid medium (NaNO_3_ 2 g L^-1^, KCl 0.5 g L^-1^, glycerol phosphate*H_2_O 0.5 g L^-1^, K_2_HPO_4_ 0.35 g L^-1^, FeSO_4_ 0.01 g L^-1^, sucrose 30 g L^-1^, agar 10 gL^-1^, adjusted to pH 6). *O. maius* and *OmΔSSP1-null* mutants were also grown in the presence of different stressor compounds. Czapek-Dox medium was supplemented with 0.3 mM Cd (as 3CdSO_4_*8H_2_O), 15 mM Zn (as ZnSO_4_*7H_2_O), 117.6 mM H_2_O_2_, 0.75 mM menadione, 0.1% (w:v) caffeic acid, 0.5% (w:v) tannic acid, 0.5% (w:v) gallic acid and 0.5% (w:v) quercetin. Prior to fungal inoculation, sterile cellophane membranes were placed on the agar surface to provide a convenient means of removing the mycelium from the plate. The membranes were first boiled for 15 min in 10 mM EDTA (disodium salt, dihydrate, SIGMA), rinsed and then autoclaved in ddH_2_O. Fungal colonies were removed after 30 days, dried over-night and weighted.

### *In vitro* mycorrhizal synthesis

Axenic *V. myrtillus* seedlings were obtained from seeds (Les Semences du Puy, Le Puy-En-Velay, France) surface sterilized in 70% ethanol (v:v) 0.2% Tween20 for 3 min, rinsed with sterile water, submerged in 0.25% sodium hypochlorite for 15 min and rinsed again with sterile water. Seeds were germinated on 1% water agar for 2 weeks in darkness before transfer to a growth chamber for 1 month.

Mycorrhiza was synthesized in petri plates containing Modified Melin-Norkrans (MMN) medium containing KH_2_PO_4_ 0.5 gL^-1^, Bovine Serum Albumin (BSA) 0.1 gL^-1^, CaCl_2_*2H_2_O 0.066 gL^-1^, NaCl 0.025 gL^-1^, MgSO_4_*7H_2_O 0.15 gL^-1^, thiamine-HCl 0.1 gL^-1^, FeCl_3_*6H_2_O 0.001 gL^-1^, agar 10 gL^-1^ and final pH 4.7. Sterile cellophane membranes, prepared as described before, were placed on the agar surface before fungal inoculation. A suspension of *O. maius* conidia in sterile deionised water was distributed on the cellophane membranes in the bottom half of the MMN petri plates. Ten germinated *V. myrtillus* seedlings were then transferred just above the conidia suspension. Plates were sealed and placed in a growth chamber (16-h photoperiod, light at 170 μmol m^−2^ s^−1^, temperatures at 23°C day and 21°C night). Roots were collected and the percentage of mycorrhization evaluated after 45 day.

As a control for the asymbiotic condition, *O. maius* was grown on the same medium used for mycorrhizal synthesis. Plates covered by cellophane membranes were inoculated with 5 mm fungal plugs and fungal colonies were removed after 45 days. Three biological replicates were prepared for each sample of the RNASeq experiment.

### RNA extraction and RNA-Seq data analyses

Total RNA was extracted and quantified from 100 mg aliquots of *O. maius* mycelium and *O. maius-*inoculated *V. myrtillus* collected 45 days after inoculation, frozen in liquid nitrogen and mechanically ground. Total RNA was extracted from *O. maius* mycelium using a Tris-HCl extraction buffer and from *V. myrtillus* mycorrhizal roots using the CTAB method, as described by Kohler et al. (2015).

Preparation of libraries from total RNA and 2 x 100bp Illumina HiSeq sequencing (RNA-Seq) was performed by IGA Technology Services (Udine, Italy). Raw reads were trimmed and aligned to the respective reference transcripts available at the JGI MycoCosm database (http://genome.jgi-psf.org/programs/fungi/index.jsf) using CLC Genomics Workbench v6. For mapping, the minimum length fraction was 0.9, the minimum similarity fraction 0.8 and the maximum number of hits for a read was set to 10. The unique and total mapped reads number for each transcript were determined, and then normalized to RPKM (Reads Per Kilobase of exon model per Million mapped reads). A summary of the aligned reads is given in Table S1. The data set was submitted to GEO (GSE63947). To identify differentially regulated transcripts in mycorrhizal tissues compared to free-living mycelium, the Baggerly’s Test (Baggerly et al. 2003) implemented in CLC Genomic workbench was used. This test compares the proportions of counts in a group of samples against those of another group of samples. The samples are given different weights depending on their sizes (total counts).

The weights are obtained by assuming a Beta distribution on the proportions in a group, and estimating these, along with the proportion of a binomial distribution, by the method of moments. The result is a weighted t-type test statistic. In addition, Benjamini & Hochberg multiple-hypothesis testing corrections with False Discovery Rate (FDR) were used. Transcripts with a more then 5-fold change and a FDR corrected p-value <0.05 were kept for further analysis.

*O. maius* Small Secreted Proteins (SSPs) were identified using a custom pipeline including SignalP v4 (1), WolfPSort (2), TMHMM, TargetP (3), and PS-Scan algorithms (4) as reported in Pellegrin et al. (2015). To assess whether symbiosis-regulated transcripts were conserved or lineage-specific (i.e., orphan genes with no similarity to known sequences in DNA databases), their protein sequences were queried against the protein repertoires of 59 fungal genomes using BLASTP with e-value 1e-5. Proteins were considered as orthologs of symbiosis-regulated transcripts pending they showed 70% coverage over the regulated sequence and at least 30% amino acid identity.

### cDNA synthesis and quantitative RT-PCR (RT-qPCR)

The expression of seven selected SSPs was evaluated by RT-qPCR. cDNA was obtained from about 1000 ng of total RNA with a reaction mix containing 10 μM random primers, 0.5 mM dNTPs, 4 μl 5× buffer, 2 μl 0.1 M DTT, and 1 μl Superscript II Reverse Transcriptase (Invitrogen) in a final volume of 20 μl. Temperature regime was: 65°C for 5 min, 25°C for 10 min, 42°C for 50 min, and 70°C for 15 min. Possible DNA contamination was tested with an additional PCR reaction using a specific DNA-primer for the *O. maius Elongation Factor1α* (*OmEF1α*) (Table S2). RT-qPCR was performed with the Rotor-Gene Q (Qiagen) apparatus. The reactions were carried out in a final volume of 15 μl with 7.5 μl of iQ SYBR Green Supermix (Bio-rad), 5.5 μl of forward and reverse primers (10 μM stock concentration; Table S2) and 2 μl of cDNA (diluited 1:10). qPCR cycling program consisted of a 10 min/95°C holding step followed by 40 cycles of two steps (15 s/95°C and 1min/60°C). The relative expression of the target transcript was measured using the 2^-ΔCt^ method (Livak & Schmittgen, 2001). The *Omβ-Tubulin* (*OmβTub*) (Table S2) was used as reference housekeeping gene. Three to five biological replicates and two technical replicates were analyzed for each condition tested. qPCR primers were designed with Primer3Plus (http://www.bioinformatics.nl/cgi-bin/primer3plus/primer3plus.cgi) and checked for specificity and secondary structure formation with PrimerBlast (http://www.ncbi.nlm.nih.gov/tools/primer-blast/) and OligoAnalyzer (eu.idtdna.com/calc/analyzer). Primers were synthesized by Eurogentec (Belgium).

### Construction of the Om*SSP1*-disruption vector and *Agrobacterium-mediated* transformation

OmSSP1-null mutants were obtained through *Agrobacterium tumefaciens*-mediated (ATM) homologous recombination. PCR reactions were used to produce the 5′ upstream flanking region (1502 bp) and the 3′ downstream flanking region (1533 bp) of the *OmSSP1* gene. PCR reactions were carried out in a final volume of 50 μl containing: 50 ng of genomic DNA of *O. maius Zn*, 1 μl dNTPs 10mM, 2.5 μl of each primer (10 μM stock concentration; Table S2), 10 μl of 5× Phusion HF Buffer and 0.5 units of Phusion Hot Start II High-Fidelity (Thermo Scientific). The PCR program was as follows: 30 s at 98°C for 1 cycle, 10 s at 98°C, 30 s at 60°C, 45 s at 72°C for 30 cycles, 10 min at 72°C for 1 cycle. Amplicons were then purified with Wizard^®^ SV gel and PCR cleanup system (PROMEGA) following manufacturer’s instructions. PCRs amplicons were cut with Xmal-HindIII (for the 5’) and BglII-HpaI (for the 3’) and cloned into the pCAMBIA0380_HYG vector (Fiorilli et al., 2016) in order to obtain the pCAMBIA0380_HYG-*ΔOmSSP1 vector* (Fig. S1). The restriction reactions were performed in a 30 μl final volume containing 0.5 μg of DNA (1 ug for the plasmid), 0.5 μl of each enzyme (from PROMEGA), 0.3 μl of BSA 100X and 3 μl of buffer 10X, overnight at 37°C. The ligase reaction was carried out in 20 μl final volume containing 50 ng of vector, 18 ng of the amplicon, 2 μl of buffer 10X and 1 μl of T4 enzyme (PROMEGA), overnight at 4°C. The obtained sequence of the vector was checked by PCRs and DNA sequencing.

The vector was cloned into *Agrobacterium tumefaciens* LBA1100, that was used to transform ungerminated *O. maius* conidia according to the protocol described in Abbà et al. (2009).

### Identification of OmSSP1-null mutants by PCR and Southern Blot

Fungal transformants were screened by PCR. A small portion of each fungal colony was collected and boiled for 15 min in 20 μl of 10 mM Tris HCl pH 8.2, vortexed for 1 min and centrifuged 15 min at room temperature. Then, 2 μl of the supernatant were used directly for PCR amplification without any other purification, using two sets of primers (Table S2). The first primer set was designed to amplify the *OmSSP1* gene (OmSSPb1r and OmSSPb1f) whereas the second set (Hyg4f e Hyg2r) was designed to amplify the portion of the *hph* gene corresponding to the Hyg-probe (Fig. S1). A OmSSP1-null mutant would yield an amplified product only with the second primer set. The putative OmSSP1-null mutants identified were validated by PCR using primers (Table S2) designed on the genome at the 5’ and 3’ of the homologous recombination site (respectively preOmSSPb1f3/Hyg6r and postOmSSPb1r3/Hyg3f). The positive OmSSP1-null mutants were further analysed through Southern hybridization analysis to verify single-copy integration of the disruption cassette in the genome. 15 μg of genomic DNA from the deletion mutants and from the wild type were digested with BamHI (PROMEGA) and size-fractionated on 1% (w:v) agarose TAE 1X gel. The separated restriction fragments were blotted onto a nylon membrane following standard procedures (see Abbà et al., 2009). Hybridization with a probe designed on the hyg-resistance cassette (Hyg-probe, Fig. S1)was performed with a chemiluminescent detection system (ECL direct DNA labelling and detection system; GE Healthcare, U.K.) was performed according to the manufacturer’s recommendations.

### Quantification of the degree of mycorrhization

To determine differences in root colonization between *O. maius* wild-type and the OmSSP1-null mutants, the percentage of mycorrhization was recorded after 1.5 months. The roots of 3 to 6 seedlings colonised by each mutant strain were collected and the whole-root system was stained overnight in a solution of lactic acid:glycerol:H_2_O (14:1:1) containing acid fuchsin 0.01% (w:v), destained twice with 80% lactic acid and observed using a Nikon Eclipse E400 optical microscope. The magnified intersections method (Villarreal-Ruiz et al., 2004) was adapted to quantify the percentage of fungal colonization of *V. myrtillus* hair roots. Roots were examined under the microscope using the rectangle around the cross-hair as intersection area at 40× magnification. A total of 60 intersections per seedling root system were scored. Counts were recorded as percentage of root colonized (RC) by the fungus using the formula: RC% = 100 ×Σ of coils counted for all the intersections/Σ of epidermal cells counted for all the intersections.

### Phylogenetic and bioinformatic analyses

The aminoacid sequences comprised between cysteine 1 and cysteine 8 of the fungal proteins listed in Table S3 were aligned using MUSCLE (Multiple Sequence Comparison by Log-Expectation; Gap Open penalty −2, Edgar, 2004) tool implemented in MEGA7 (Tamura et al., 2007). Maximum likelihood analysis (Guindon & Gascuel 2003) was conducted using www.phylogeny.fr in advanced mode (Dereeper et al., 2008). The phylogenetic tree was reconstructed using the maximum likelihood method implemented in the PhyML program (v3.1/3.0 aLRT).

Bioinformatic analyses of protein primary sequences were performed using on line tools. Blastp searches on the Uniprot database (The UniProt Consortium, 2017) identified the closest protein matches. Hydropathy profiles were generated with ProtScale tools (http://web.expasy.org/protscale/), while the grand average of hydropathicity (GRAVY), the predicted amino acid number and molecular weight were calculated using the ProtParam tool (http://web.expasy.org/protparam/). The intrinsic solubility profiles were obtained with the camsolintrinsic calculator (http://www-mvsoftware.ch.cam.ac.uk/index.php/camsolintrinsic). The representation of the residues hydrophobicity of the aligned sequences of each clade was obtained using http://www.ibi.vu.nl (Simossis & Heringa 2005).

### The yeast signal sequence trap assay

Functional validation of the predicted signal peptide of OmSSP1 was conducted with a yeast signal sequence trap assay (Plett et al., 2011). The pSUC2-GW gateway vector carries a truncated invertase (SUC2) lacking both its initiation methionine and signal peptide. cDNA encoding the predicted OmSSP1 signal peptide was cloned into pSUC2-GW plasmids using BP/LR technologies (Invitrogen). Then, yeast cells (YTK12 strain) were transformed with 200 ng of the individual pSUC2-GW/OmSSP1 plasmid using the lithium acetate method (Gietz & Schiestl, 2007). Transformants were grown on yeast minimal medium with SD/W^-^ medium (adenine hemisulfate 40 mgL^-1^, L-uracil 40 mgL^-1^, L-histidine hydrochloride 40 mgL^-1^, L-leucine 40 mgL^-1^). After the autoclave 200 ml of SD 5x solution containing 3.5 gL^-1^ of drop out mix and 33.5 gL^-1^ of yeast nitrogen base without amino acids (adjusted to pH 5.6) and glucose 20 gL ^-1^ were added to the SD/W^-^ medium. To assay for invertase secretion, colonies were grown overnight at 30°C with shaking 200 rpm and diluted to an OD_600_ = 1, then 5 μl of serial dilution of the yeast culture were plated onto YPSA medium containing sucrose (10 gL^-1^ yeast extract, 20 gL^-1^ peptone, 20 gL^-1^ agar amended with 2 gL^-1^ sucrose and 60 μg mL^-1^ antimycin A after autoclaving, pH 6.5).

### Statistical analyses

The significance of differences among the different treatments was statistically evaluated by ANOVA with Tukey’s pairwise comparison as post-hoc test for multiple comparisons for normal distributed data. Statistical elaborations of growth and biomass data were performed using PAST statistical package, version 2.17 (Hammer et al., 2001). The differences were considered significant at a probability level of *p*<0.05.

## Results

### The *O. maius* genome contains several SSPs up-regulated during mycorrhizal symbiosis with *V. myrtillus*

Among the 16,703 genes found in the *O. maius* genome (Kohler et al., 2015), 445 genes (~38% of the total *O. maius* predicted secretome) code for putatively secreted proteins smaller than 300 amino acids (Table S4a). The transcriptomic analysis of *O. maius* under free-living conditions (FLM) and in symbiosis with *V. myrtillus* (MYC) indicated that about 24% (278/1163) of the genes coding for putatively secreted proteins were induced in symbiosis (Fold Change>5, p-value <0.05), 90 of them corresponding to SSPs. Many of these mycorrhiza-induced SSPs (MiSSP) were strongly up-regulated in symbiosis (13 with FC≥400) or mycorrhiza-specific (Table S4b). Only 32 were cysteine enriched (C>3%), a feature normally attributed to SSPs (Kim et al., 2016). About half (49/90) contained PFAM motifs specific of CAZymes (especially glycoside hydrolases, GHs), lipases, hydrophobins and peptidases, whereas the remaining 41 contained no known PFAM domain. None of these MiSSPs contained a nuclear localization signal motif and 6 of them featured a KR rich sequence, i.e. a motif characterized by basic aminoacids (like lysine and arginine) supporting the plant nucleus entry (Table S4b).

Genes orthologous to the *O. maius* MiSSPs were identified by genomic comparative analyses with 59 taxonomically and ecologically distinct fungi (Table S4b), including three other ERM fungi in the Leotiomycetes *(Meliniomyces bicolor, M. variabilis* and *Rhizoscyphus ericae)* belonging to the “*R. ericae*” aggregate (Vrålstad et al., 2000). Ten of the 90 *O. maius* symbiosis-induced SSPs resulted to be specific for *O. maius*, whereas 2 to 834 orthologous genes were found for the other 80 SSPs (Table S4b). Many *O. maius* SSPs orthologs were found in the other three ERM fungal species, although no ERM specific SSPs could be identified. The highest number of *O. maius* SSPs orthologs was found in the genomes of pathogenic and saprotrophic fungi (64 and 73 respectively), whereas only 38 orthologous genes were found in the genomes of 12 ECM fungi (Table S4b).

We selected seven *O. maius* SSPs for further analyses, with a preference for those uniquely or highly expressed in symbiosis that did not contain PFAM motifs with known functions (Table 1). Three different software (PrediSi, SIgnalP 4.1, Phobius) confirmed the presence of a signal peptide (Table 1), but alignment of the primary sequences indicated very low similarities. Cysteine enrichment >3% was only observed for OmSSP1 (8.6%), the most highly symbiosis-induced SSP (Table 1). A positive GRAVY value was found for OmSSP1 (0.53) and OmSSP3 (0.40), which indicates hydrophobic protein regions, whereas the negative values found for the other five SSPs indicate overall protein hydrophilicity (Table 1). The expression of these OmSSPs was investigated in free-living and in mycorrhizal *O. maius* by RT-qPCR experiments, which confirmed a significant up-regulation of all selected SSP genes in symbiosis, and in particular a very strong induction of OmSSP1 (Fig. 1).

**Figure 1.**
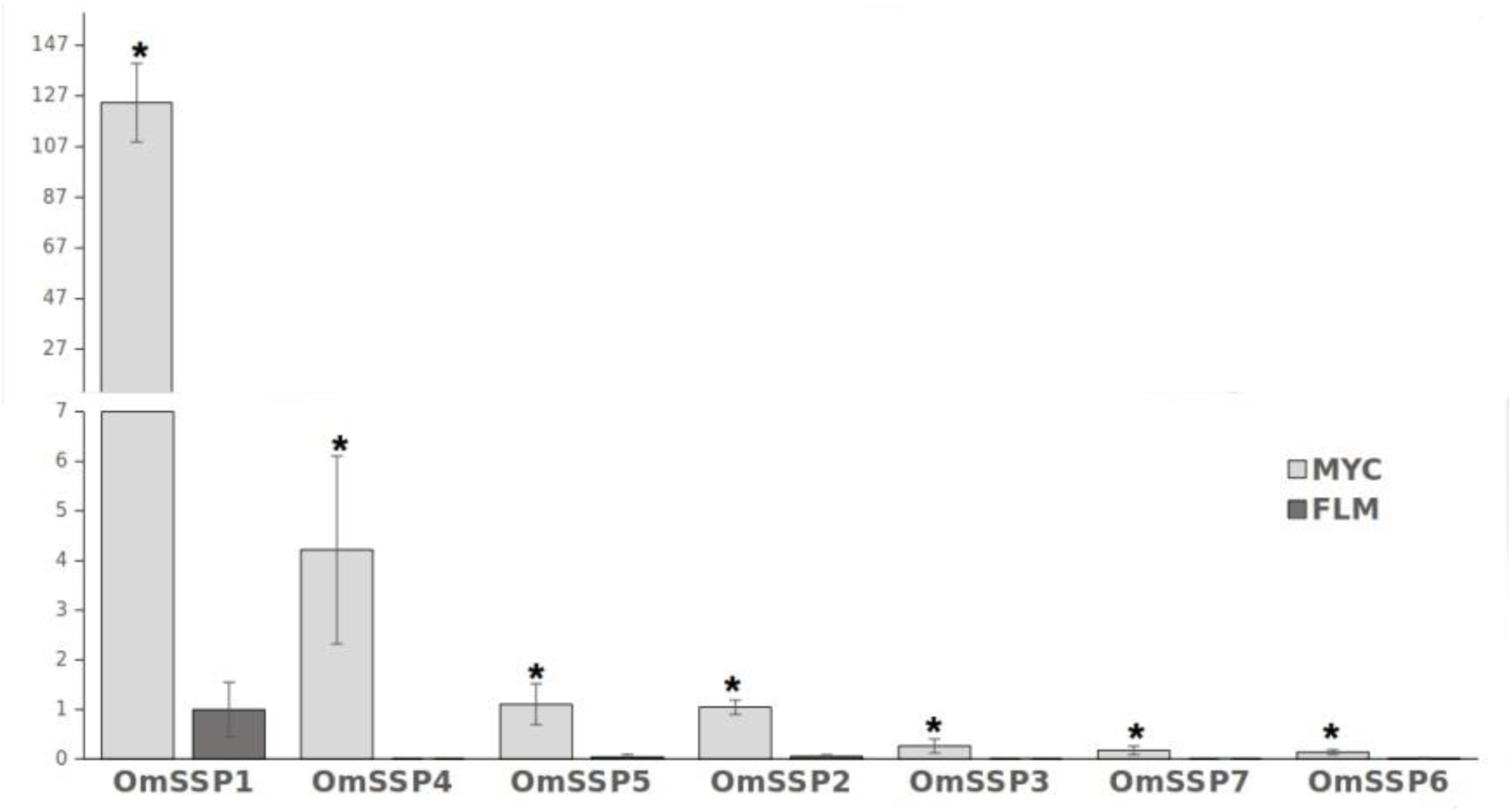
RT-qPCR validation of selected *O. maius* SSPs expression level in mycorrhizal tissues of *V. myrtillus* (MYC) as compared to the free living mycelium (FLM). Relative expression (mean of ΔCt values) of 7 *O. maius* SSPs normalized to *OmTub* transcripts. Bars represent the mean ±SD. Asterisks (*) indicates a statistically significant difference (*p* < 0.05) (ANOVA, Tukey’s post hoc test).

**Table 1.** Expression level, biochemical and bioinformatic features of seven selected *O*. maius mycorrhizal-induced SSPs.

| Name |  | Expression |  |  |  | Biochemical features |  |  |  |  | N. of orthologs<br>(see also Table S4c) |  |
| --- | --- | --- | --- | --- | --- | --- | --- | --- | --- | --- | --- | --- |
| Selected SSPs | Protein ID | FLM Means | MYC Means | Fold Change | p-value | Predicted domains, repeats PFAM motifs | C residues (%) | GRAVY index | Protein length | Molecular weight (kDa) | *O. maius* | Total (60 genomes) |
| **OmSSP1** | 182936 | 0.685 | 13734 | 20056 | 5.07 E-20 | 1 LCR, NO PFAM | 8 (8.60) | + 0.53 | 93 | 9.2 | 1 | 4 |
| **OmSSP2** | 160794 | 0.103 | 409 | 3961 | 8.84 E-44 | 1 LCR, PFAM: DUF4360 | 4 (1.58) | − 0.29 | 253 | 27.6 | 3 | 82 |
| **OmSSP3** | 56093 | 0.044 | 28 | 628 | 2.96 E-10 | NO PFAM | 1 (0.54) | + 0.40 | 188 | 19.7 | 1 | 1 |
| **OmSSP4** | 20160 | 0 | 105 | MYC | 7.56 E-10 | 2 internal repeats, NO PFAM | 0 (0.00) | − 0.48 | 102 | 11.3 | 1 | 1 |
| **OmSSP5** | 23797 | 0 | 48 | MYC | 1.55 E-06 | 2 internal repeats, NO PFAM | 0 (0.00) | − 0.24 | 108 | 11.8 | 1 | 1 |
| **OmSSP6** | 125454 | 0 | 34 | MYC | 1.50 E-22 | NO PFAM | 2 (0.91) | − 0.06 | 220 | 23.0 | 2 | 14 |
| **OmSSP7** | 25811 | 0 | 16 | MYC | 0.003 | 1 LCR, NO PFAM | 6 (2.37) | − 0.19 | 253 | 26.5 | 3 | 172 |

LCR: Low complexity region
MYC: Transcripts could not be identified in the free living mycelium (FLM) and were considered as mycorrhiza specific

### OmSSP1 shows molecular similarities with fungal proteins annotated as hydrophobins

*OmSSP1* is a single copy gene located in the scaffold 14 of the *O. maius* genome. The coding region contains 354 nucleotides, with 2 exons and 1 intron. In addition to the signal peptide, OmSSP1 is rich in glycine (16.1%) and leucine (12.9%) and contains 8 cysteine residues (Fig. 2). The intrinsic calculated solubility and solvent accessibility highlighted at least three poorly water soluble regions. The majority of hydrophobic residues in OmSSP1 are clustered between amino acid residues 41-56 and 79-93 (Fig. 2). OmSSP1 secondary structure predictions indicated a disordered folded state for 46% of the protein, partly due to the presence of a low complexity region (LCR) before the first cysteine (Fig. 2), whereas 16% and 20% of the protein structure could form helix and beta-sheet structures, respectively (not shown).

**Figure 2.**
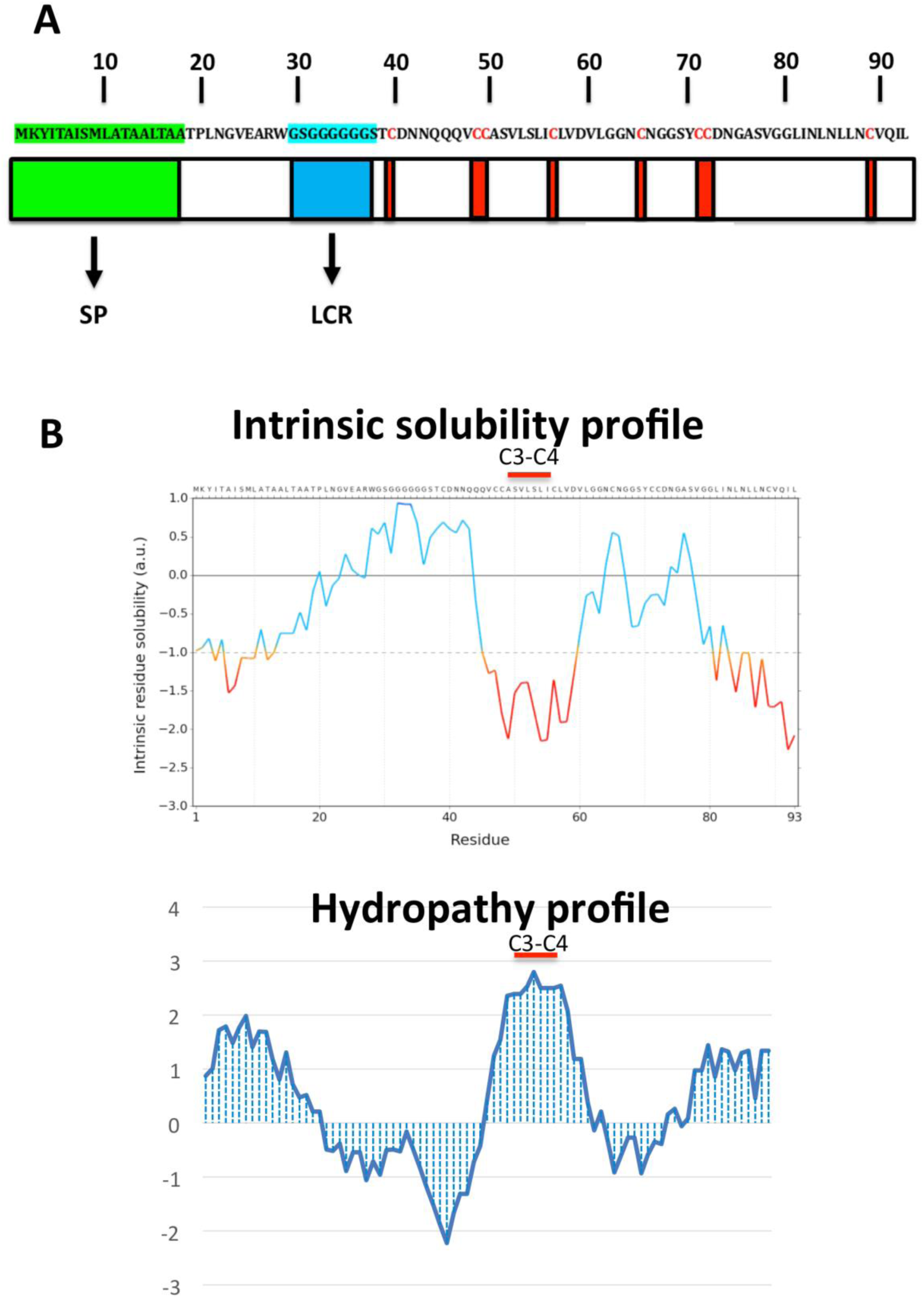
OmSSP1 structural and biochemical features predicted through bioinformatics tools. (**A**) Schematic representation of OmSSP1 primary amino acid sequence. The green bar represents the signal peptide (SP), the blue bar the low complexity region (LCR) and the red bars the cysteine (C) residues. (**B**) Calculation of the intrinsic solubility profile (ISP) and hydrophaty profile (HP). For the ISP, scores larger than 1 indicate highly soluble regions, while scores smaller than −1 indicate poorly soluble regions. For the HP, hydrophobic aa show positive peaks with values above 0 whereas hydrophilic aa show negative peaks.

Although no known domains could be found in the predicted OmSSP1 protein sequence by PFAM database searching, nor a functional classification in the InterPro database, BlastP searches both in the UniProt and in the RefSeq databases yielded, as best identified protein matches, fungal hydrophobins or hydrophobin-like proteins (Table 2). Therefore, we compared the protein sequence of OmSSP1 with those of the four annotated hydrophobins in *O. maius* and with other fungal hydrophobins (Linder et al., 2005; Seidl-Seiboth et al., 2011, Grigoriev et al., 2014). Three out of the four *O. maius* hydrophobins showed a GRAVY score above 0.6, indicating an overall hydrophobicity higher than OmSSP1 (0.53), whereas the GRAVY score for *O. maius* hydrophobin 4 was only 0.32 (Table S5).

**Table 2.**
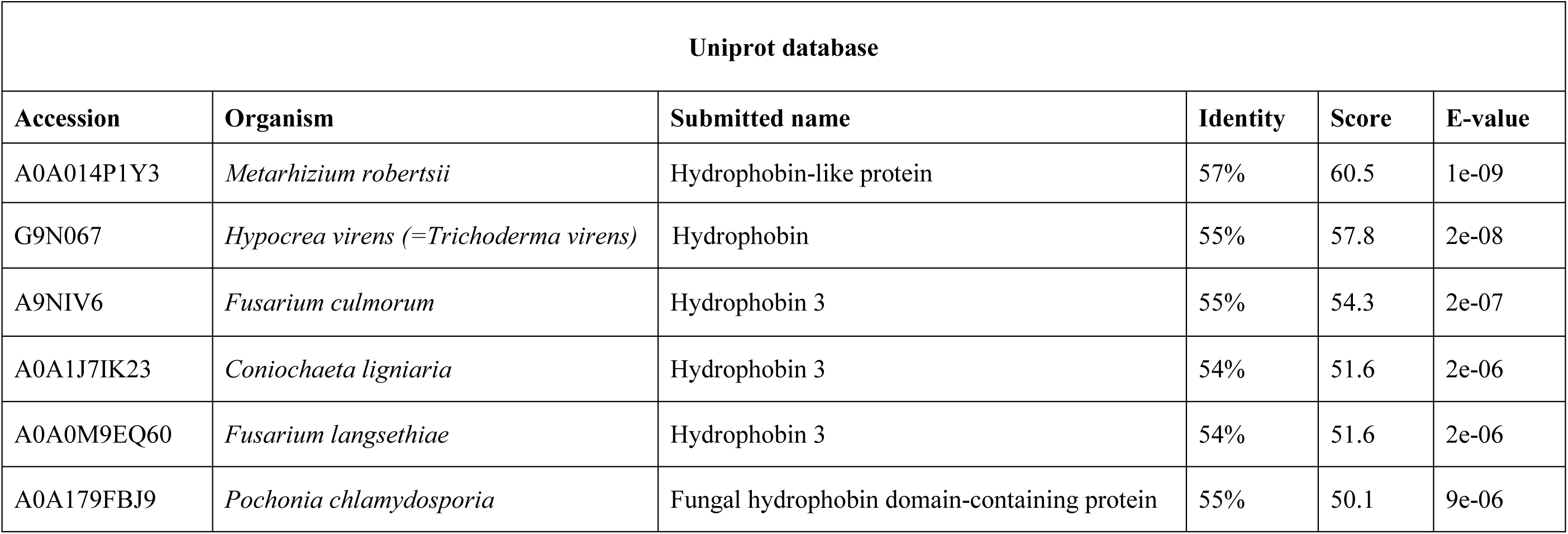
Results of BlastP searches on the Uniprot database using the OmSSP sequence as query

| Uniprot database |  |  |  |  |  |
| --- | --- | --- | --- | --- | --- |
| Accession | Organism | Submitted name | Identity | Score | E-value |
| A0A014P1Y3 | *Metarhizium robertsii* | Hydrophobin-like protein | 57% | 60.5 | 1e-09 |
| G9N067 | *Hypocrea virens* (= *Trichoderma virens*) | Hydrophobin | 55% | 57.8 | 2e-08 |
| A9NIV6 | *Fusarium culmorum* | Hydrophobin 3 | 55% | 54.3 | 2e-07 |
| A0A1J7IK23 | *Coniochaeta ligniaria* | Hydrophobin 3 | 54% | 51.6 | 2e-06 |
| A0A0M9EQ60 | *Fusarium langsethiae* | Hydrophobin 3 | 54% | 51.6 | 2e-06 |
| A0A179FBJ9 | *Pochonia chlamydosporia* | Fungal hydrophobin domain-containing protein | 55% | 50.1 | 9e-06 |

Two classes of fungal hydrophobins have been distinguished by Wessel (1994) on the basis of the hydropathy profile and of slightly different motifs between the eight positionally conserved cysteine residues, Class I (C-X_5–8_-CC-X_17–39_-C-X_8–23_-C-X_5–6_-CC-X_6–18_-C-X_2–13_) and Class II (C-X_9–10_-CC-X_11_-C-X_16_-C-X_8–9_-CC-X_10_-C-X_6–7_). OmSSP1 bears a signature motif similar to class I hydrophobins (C-X_7_-CC-X_7_-C-X_8_-C-X_5_-CC-X_16_-C-X_2_), with the exception of a shorter sequence (only 7 aminoacids) between cysteines 3 and 4, expected to host the most hydrophobic protein region. Although shorter, the C_3_-C_4_ loop of OmSSP1 showed a conserved hydrophobic core composed by valine (V) and leucine (L), thus explaining the negative values in solubility profile and positive values in hydropathy profile (Fig. 2). Hydrophobins are amphiphilic molecules (Whiteford & Spanu, 2002; Rineau et al., 2017), and OmSSP1 has a very hydrophilic stretch rich in G before the C_3_-C_4_ loop, corresponding to the LCR (Fig. 2).

To better understand the phylogenetic relatedness of OmSSP1 with fungal hydrophobins, we aligned the OmSSP1 protein sequence with the four annotated *O. maius* hydrophobins and with Class I and Class II annotated hydrophobins from Ascomycetes (listed in Table S3). A phylogenetic tree built on the complete C_1_-C_8_ sequence alignment is shown in Fig. 3. However, since the different proteins showed a highly variable sequence length of the C_3_-C_4_ loop, ranging from 4 to 39 amino acids, a more conserved phylogenetic tree was generated by Maximum Likelihood (ML) without this protein region, to avoid possible bias due to the different protein lengths (Fig. S2). In both trees, class II hydrophobins and two of the four *O. maius* hydrophobins (Oidma2 and Oidma3) grouped in a single, well supported cluster. The two other *O. maius* hydrophobins grouped in a well-supported cluster together with characterised Class I hydrophobins (Figs. 3 and S2). Although the position of some fungal proteins differed in the two ML trees, most terminal clades (A to D) were maintained and well supported. OmSSP1 clustered in Clade C (Figs. 3 and S2) together with other proteins reported as hydrophobins (Table S3) and featuring a short C_3_-C_4_ loop (X_4-9_). The complete C_1_-C_8_ sequence alignment of proteins in Clade C is shown in Fig. 4. As in OmSSP1, most amino acids in the short C_3_-C_4_ loop of other proteins in this clade were hydrophobic. Clade B (Figs. 3 and S2) was another well-supported clade containing proteins also featuring a very short (X_8-9_) C_3_-C_4_ loop (Fig. 4). Clade B included a *Trichoderma atroviride* hydrophobin (Triat1) described by Seidl-Seiboth et al. (2011) as a member of a novel subclass in Class I hydrophobins. Unlike Clade C, proteins in Clade B featured mainly hydrophilic amino acids in the C_3_-C_4_ loop (Fig. 4). Clade A included the two *O. maius* hydrophobins Oidma1 and Oidma4.

**Figure 3.**
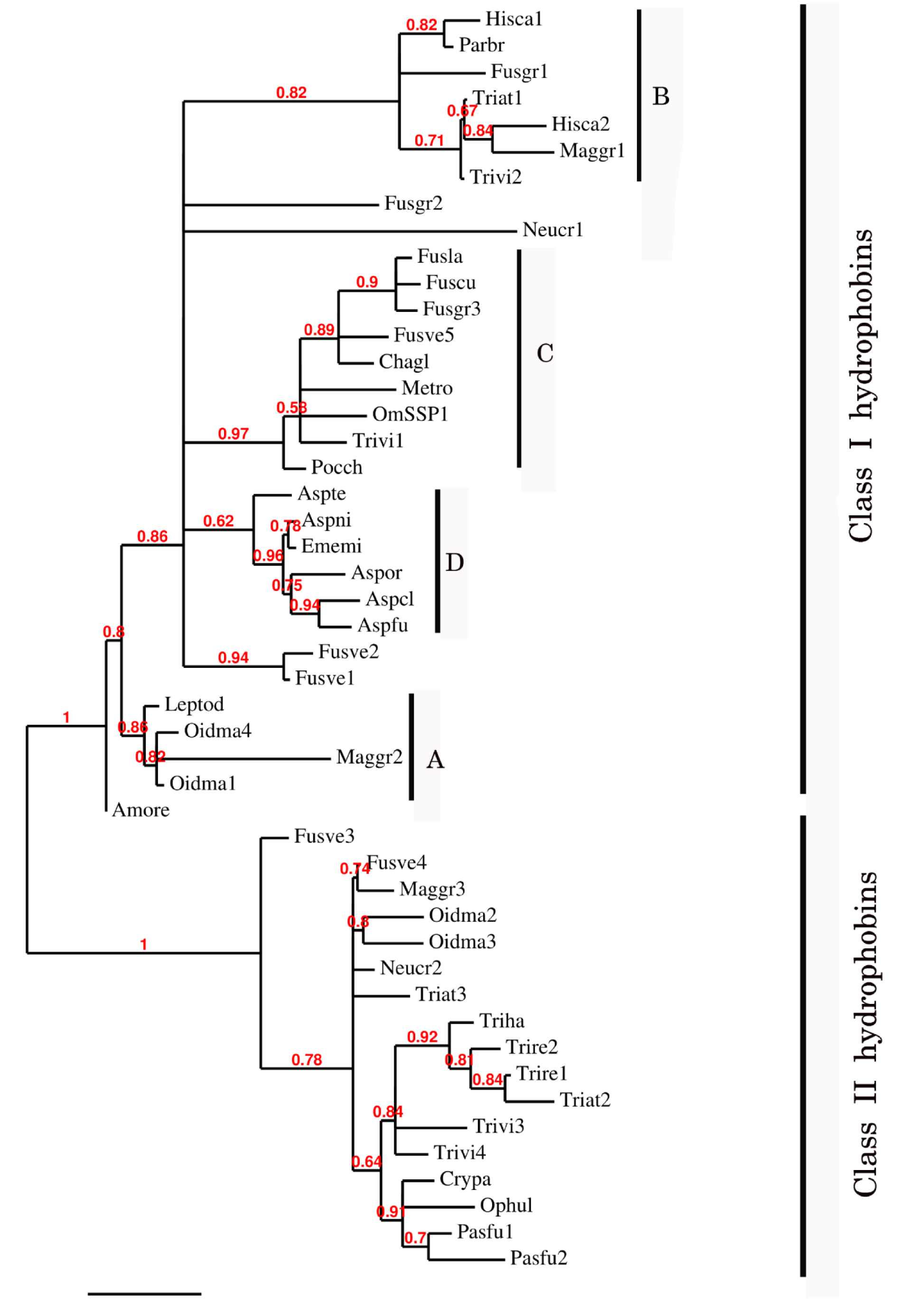
Phylogenetic tree of OmSSP1 and *O. maius* hydrophobins with other annotated hydrophobins from Ascomycetes. The analysis included protein sequences annotated or described as Ascomycetes class I and class II hydrophobins (listed in Table S3), OmSSP1 and the four *O. maius* hydrophobins. This sequence alignment considered the complete amino acid sequence comprise between C1 and C8. Muscle algorithm implemented in MEGA7 (Tamura et al., 2007) was used to generate the multiple protein sequence alignment. The phylogenetic tree was reconstructed on the Phylogeny.fr platform (Dereeper et al., 2008) using the maximum likelihood method (Guindon & Gascuel 2003) implemented in the PhyML program (v3.1/3.0 aLRT). The WAG substitution model was selected assuming an estimated proportion of invariant sites (of 0.088) and 4 gamma-distributed rate categories to account for rate heterogeneity across sites. The gamma shape parameter was estimated directly from the data (gamma=7.097). Reliability for internal branch was assessed using the aLRT test (SH-Like). Graphical representation and edition of the phylogenetic tree were performedwit TreeDyn (v198.3; Chevenet et al., 2006).

**Figure 4.**
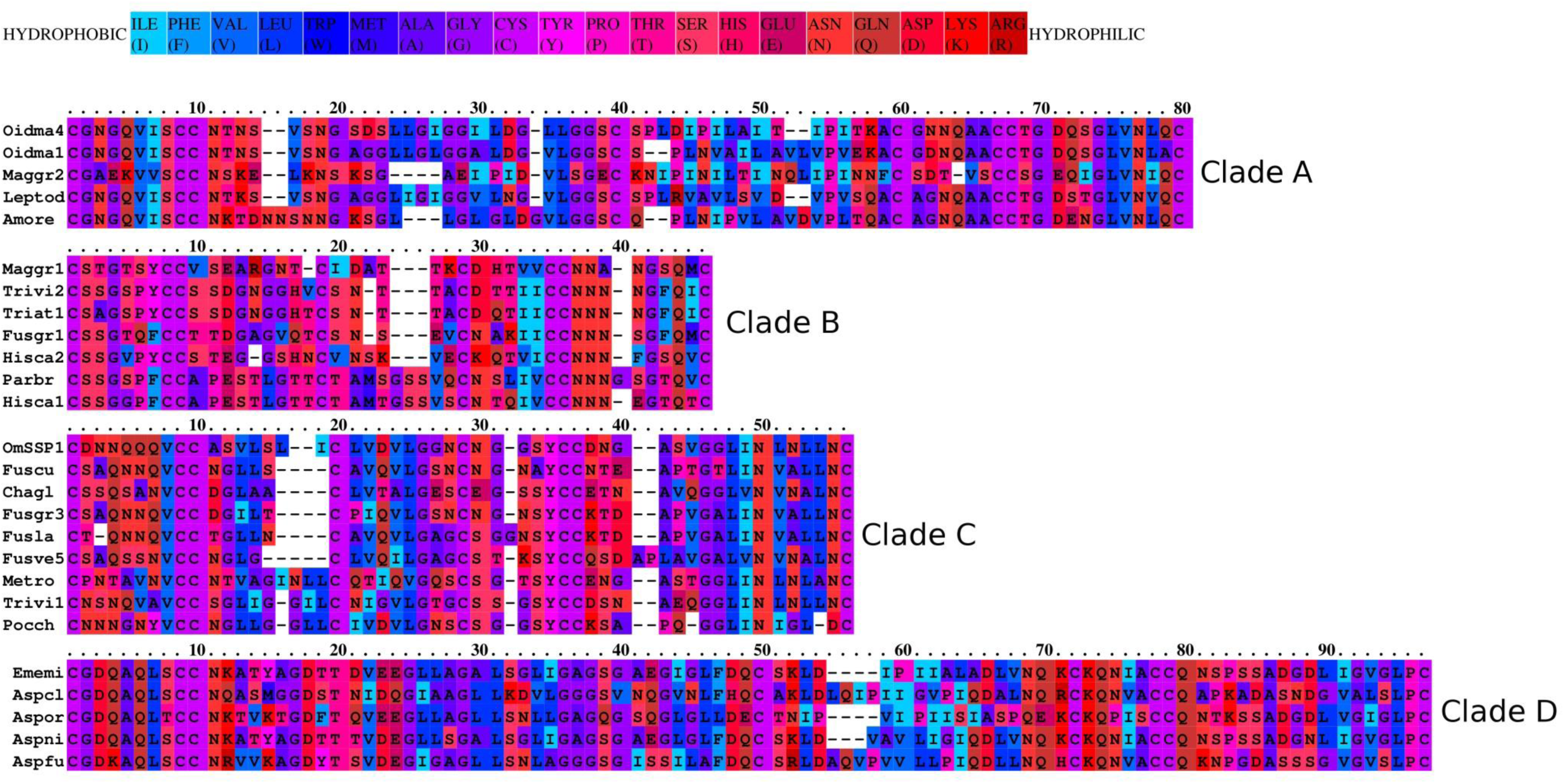
Aminoacid hydrophobicity properties of the aligned proteins. The protein sequences belonging to the four clades observed in Fig.3 were aligned by using the Praline tool of the IBIVU server (http://www.ibi.vu.nl/programs/pralinewww/; Simossis et al., 2005). The hydrophobicity scale used is from Eisenberg et. al. (1984).

### The yeast invertase secretion assay indicates that OmSSP1 is secreted

The predicted OmSSP1 signal peptide was functionally validated in a yeast signal sequence trap assay. This test is based on the yeast requirement for a secreted invertase (SUC) to grow on sucrose amended media (Klein et al., 1996). The pSUC-GW (Jacobs et al., 1997) gateway vector carries a truncated invertase that lacks its signal peptide (SUC2). The cDNA coding for the putative OmSSP1 signal peptide (OmSSP1_SP) was fused in frame to the yeast SUC2 invertase, and the recombinant pSUC-GW vector was transformed into the invertase-deficient yeast strain YTK12. As positive control, the MiSSP7 (Plett et al., 2011) signal peptide (MiSSP7_SP) and the yeast wild type signal peptide (SUC2_SP^+^) were used, whereas the empty vector was used as negative (mock) control. All transformants grew on SD-W and YPGA control media containing glucose, whereas only OmSSP1_SP and the two positive controls rescued YTK12 yeast’s growth on YPSA media containing sucrose, indicating that OmSSP1 contains a secretion sequence that is functional in yeast (Fig. 5).

**Figure 5.**
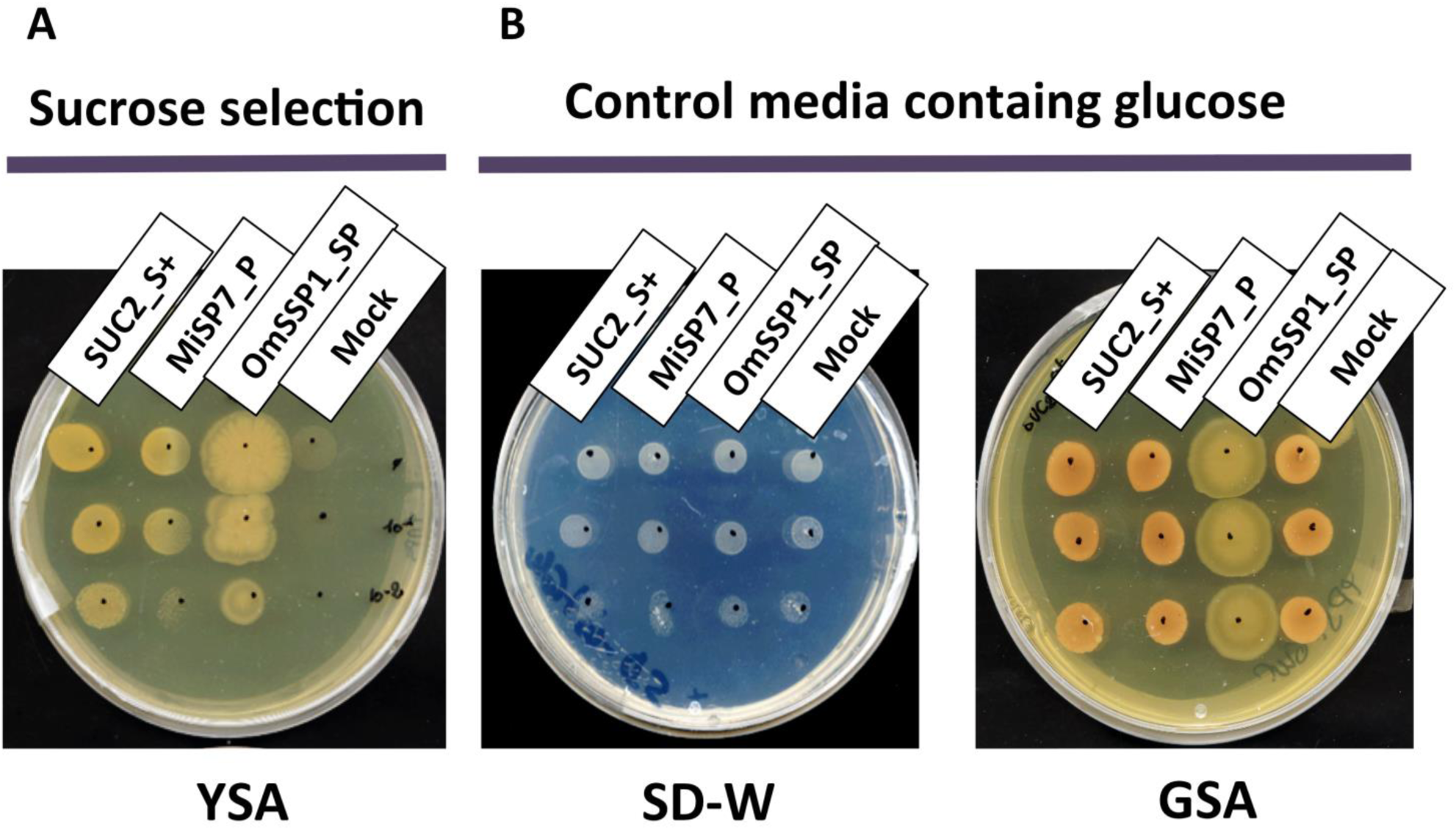
OmSSP1 contains a secretion sequence that is functional in yeast. (**A**) OmSSP1 signal peptide: upon sucrose selection the OmSSP1 signal peptide (OmSSP1_SP) rescues the functionality of the yeast invertase as well as of the other two positive controls (MiSSP7 signal peptide - MiSSP7_SP - and the wild type sequence of the yeast signal peptide - SUC2_SP^+^). The empty vector (mock) was used as a negative control. (**B**) Control media (SD-W and YPGA) containing glucose restore the growth ability of all yeast transformants.

### Growth of *OmΔSSPl* mutants is not impaired under stressful conditions, but they have a reduced capability to colonize *V. myrtillus* roots

*O. maius* can be genetically transformed and gene disruption can be obtained by homologous recombination (Martino et al., 2007; Abbà et al., 2009). To investigate the biological function of OmSSP1 in *O. maius,* knock-out mutants *(OmΔSSP1)* were obtained through AMT transformation using a vector containing the hpd-cassette (Fig. S1). Hygromycin-resistant colonies were screened by PCR, and eight putative homologous recombinants out of 742 screened fungal transformants could be identified. Southern blot hybridization with a probe to the Hygromycin cassette showed a single band of the expected size for four out of the eight candidate mutants (Fig S3). To further confirm the vector insertion site, PCR amplifications were performed with primers designed to amplify the genome regions flanking the inserted pCAMBIA0380*ΔOmSSP1* disruption cassette (Fig. S1, Table S2), followed by sequencing of the amplicons. Three *O. maius* transformants (*OmΔSSP1*^150^, *OmΔSSP1*^377^, *OmΔSSP1*^412^) were confirmed as deletion mutants for the *OmSSP1* gene. These three OmSSP1-null mutants were not affected in mycelium morphology or growth rate when inoculated on Czapek-Dox solid medium (not shown). OmSSP1 deletion did not modify the wettability phenotype of the *O. maius* mycelium (Fig. S4), but it should be noted that OmSSP1 gene expression was very low in the FLM.

It has been recently suggested that, in FLM, SSPs may increase fungal tolerance to toxic compounds (such as aromatic compounds or reactive oxygen species) released during substrate degradation (Valette et al., 2017). We therefore investigated whether deletion of the OmSSP1 gene reduced *O. maius* fitness when exposed to different stress inducers. As shown in Fig. S5, growth of the three *OmΔSSP1* mutants was not significantly affected by any of the stress conditions tested, namely two toxic heavy metals (Cd and Zn), molecules causing oxidative stress (H_2_O_2_ and menadione) and plant-derived organic compounds displaying toxic/antimicrobical effects (caffeine, tannic acid, gallic acid, quercetin and caffeic acid).

We then investigated the symbiotic ability of the three *OmΔSSP1* mutants on seedlings of the host plant *V. myrtillus* as compared with the wild type *O. maius* strain (Fig. 6). Although there were no statistically significant differences in plant biomass after 45 days of co-culture, a statistically significant (p<0.05) reduction in the percentage of root colonization was measured for all *OmΔSSP1* mutants, when compared with the wild type *O. maius* strain (Fig. 6).

**Figure 6.**
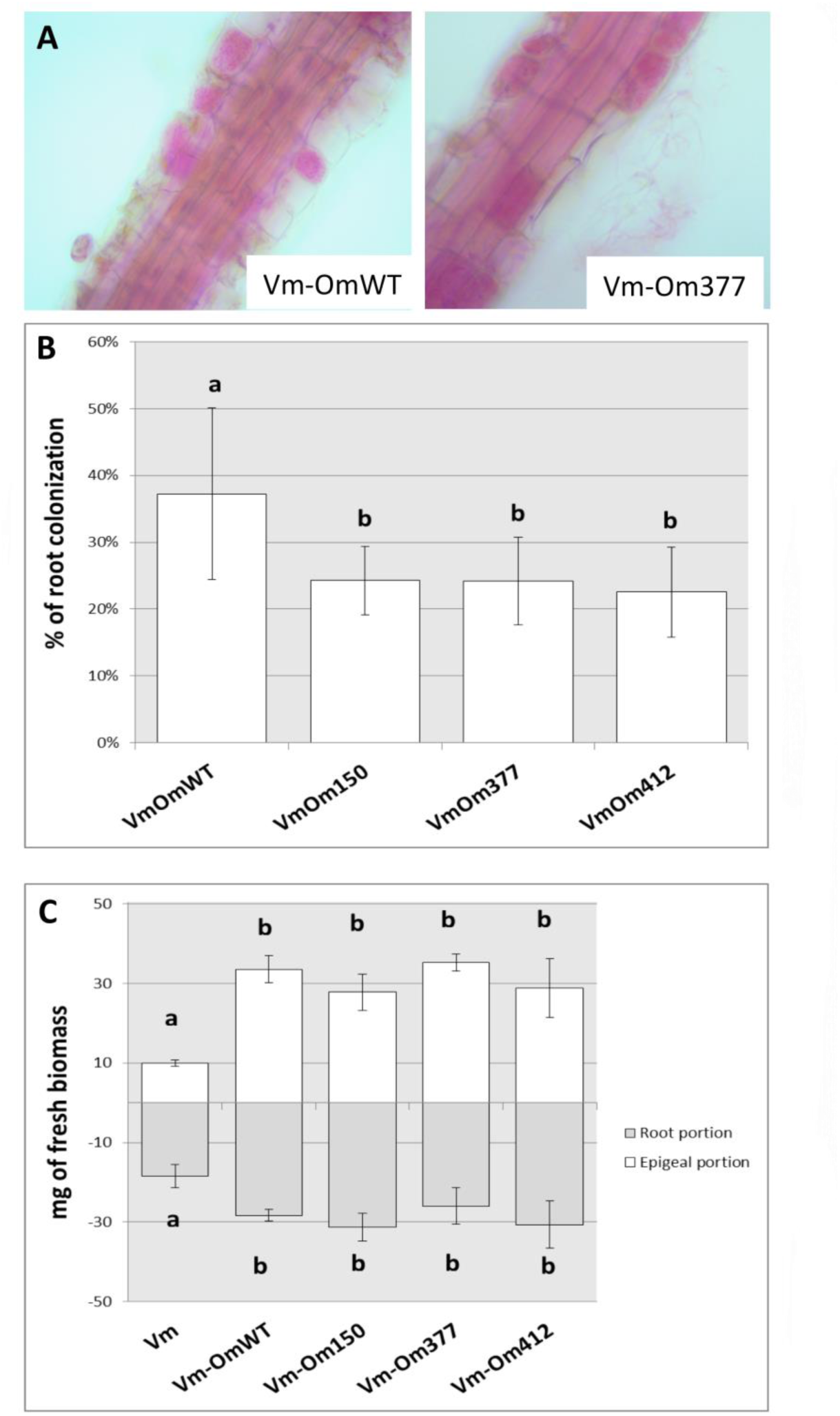
OmΔSSP1 mutants have a reduced capability to colonize *V. myrtillus* roots. Roots of *V. myrtillus* observed after 1.5 months of co-culture with *O. maius* WT and with three OmSSP1 null-mutants strains: (A) Hyphal coils fuchsine-stained were observed in *V. myrtillus* roots using the light microscope. (B) The percentage of the root colonization was significantly lower for the OmSSP1 null-mutants as compared to the *O. maius* WT. (C) Quantification of fresh plant biomass (roots - grey bars - and aboveground portions – white bars) of *V. myrtillus* plants grown alone, in the presence of the *O. maius* WT strain or of the OmSSP1 null-mutants. All pictures were taken at the same magnification. Bars represent the mean ±SD, n=5 (each biological replicate represents the total biomass of eight *V. myrtillus* seedlings grown in an individual plate). Different letters indicate statistically significant difference (*p* < 0.05) (ANOVA, Tukey’s post hoc test).

## Discussion

### Similar to other mycorrhizal fungi, the ERM fungus *O. maius* has a wide array of SSPs

Effector-like SSPs have been found to be secreted by ECM and AM fungi and to be instrumental for plant colonisation (Plett & Martin, 2015; Martin et al., 2016). SSPs are also encoded in the genome of the model ERM fungus *O. maius*. Overall, the *O. maius* genome contains 445 SSPs, corresponding to 2.6% of the total number of *O. maius* genes. Similar percentages were reported for the ECM fungus *L. bicolor* (Pellegrin et al., 2015) and for saprotrophic fungi (Valette et al., 2017). The 90 symbiosis-induced SSPs correspond to about 20% of the total *O. maius* SSPs, a percentage similar to the ECM fungus *L. bicolor* (Kohler et al., 2015) and the AM fungus *R. irregularis* (Tisserant et al., 2013).

In *O. maius*, the most highly up-regulated SSP (OmSSP1) was 20,000 times more expressed in mycorrhizal roots than in the FLM. Bioinformatic analysis of *O. maius* symbiosis-induced SSPs revealed that 45.5% (41/90) correspond to orphans genes with no known PFAM domains. Although many fungal effector SSPs are targeted to the host plant nucleus (Lo Presti et al., 2015), none of the *O. maius* symbiosis-induced SSPs showed features supporting a localization in the plant nucleus. Species specific SSPs (SSSPs), defined as SSPs with no homology in other species, have been found in both AM (Salvioli et al., 2016; Tang et al., 2016; Sędzielewska Toro & Brachmann, 2016) and ECM fungi (Pellegrin et al., 2015) and they are considered to be likely involved in the promotion of host-specific interactions (Pellegrin et al., 2015). Comparative genomics revealed that 27% of the total *O. maius* SSPs are *O. maius* specific (Table S5a). This value falls in the proportion of species-specific SSPs (SSSPs) for symbiotic organisms (25-50%) predicted by Kim et al. (2016), although only 10 out of the 90 *O. maius* symbiosis-induced SSPs were species-specific.

Whereas lifestyle-specific SSPs have been found for ECM fungi (Pellegrin et al., 2015), no ERM specific SSPs (i.e. shared by the four ERM fungi and with no orthologous in the other fungi used for comparative analysis) could be found. ERM and ECM fungi also differed because the comparative analysis showed that the highest number of *O. maius* symbiosis-induced SSPs orthologous belong to pathogenic and saprotrophic fungi, whereas ECM fungi share most of their SSPs with saprotrophs such as brown rot, white rot, and litter decayers (Pellegrin et al., 2015).

### OmSSP1, the most highly expressed *O. maius* MiSSP, may be a distinctive type of hydrophobin

Hydrophobins are small secreted proteins less than 200 amino acids long, with a secretion signal and a pattern of eight cysteine residues recurring in the sequence (Whiteford & Spanu, 2002). A unique three-dimensional folding comes from these features, keeping exposed the hydrophobic residues and rendering them amphiphilic (Rineau et al., 2017). Two classes of hydrophobins have been recognised (Wessels 1994): Class I, with higher sequence variability and more stable superstructures, is found in both Asco- and Basidiomycetes, while Class II has only been found in Ascomycetes (Kershaw & Talbot, 1998). Hydrophobins are abundantly expressed during fungal development, pathogenesis and symbiosis (Wösten, 2001; Whiteford & Spanu, 2002). Being amphiphilic, they could behave as biosurfactants and facilitate fungal adhesion to organic matter and its decomposition (Rineau et al., 2017). Hydrophobins are also instrumental for fungal hyphae to form aerial structures and to adhere to each other and/or to hydrophobic surfaces, such as the plant leaf surface during pathogenesis. Symbiosis-upregulated hydrophobins have been found in the ECM fungi *Pisolithus tinctorius* (Tagu et al., 2001) and in *L. bicolor* (Martin et al., 2008; Plett et al., 2012), where they could play a role in establishing hyphal aggregation in the symbiotic interfaces (Raudaskoski & Kothe, 2015).

The *O. maius* genome features four annotated hydrophobins containing the PFAM and InterPro hydrophobin domains. Rineau et al. (2017) suggested that all *O. maius* hydrophobins belong to Class I, but our phylogenetic analysis (that also included Class II hydrophobins) showed that two proteins (Oidma1 and Oidma4) belong to Class I and two (Oidma2 and Oidma3) to Class II hydrophobins. OmSSP1 shares some features with Class I hydrophobins and clusters with annotated hydrophobins in this Class, but it was not identified as a hydrophobin because it lacks the corresponding PFAM and InterPro domains, possibly because of the shorter C3-C4 region.

Amino acid features, such as charge and hydrophobicity, can influence hydrophobin structure and function. Thus, the amino acidic composition of the C_3_-C_4_ loop as well as of the N-terminal region of hydrophobins may influence the wettability and the substrate-attachment preference of the protein (Linder et al., 2005; Kwan et al., 2006). In this respect, it is interesting to note that proteins in Clade B and Clade C, both showing a C_3_-C_4_ loop unusually short for Class I hydrophobins, feature amino acid sequences with very different hydrophobicity (Fig.4), suggesting they may represent structurally and functionally diverse subclasses of Class I hydrophobins. The low complexity region found in OmSSP1 is also unusual for hydrophobins and could be considered a recently evolved trait of this protein (Toll-Riera et al., 2012). Its presence suggests for OmSSP1 a low propension to aggregate and to form alpha-helices and beta-sheets, three properties often correlated with the ability of hydrophobins to pile up in needle-like (amyloid) structures (Rineau et al., 2017).

A high number of hydrophilic residues (asparagine especially) were found in the N-amino terminal region of OmSSP1. According to Linder et al. (2005), the amino terminal region of hydrophobins could have important roles in the specific function of individual proteins. For example, hydrophobins featuring high number of exposed hydrophilic residues at the N-terminal region were found to be overexpressed in mycorrhizal tissues (Whiteford & Spanu, 2002; Rineau et al., 2017).

### OmSSP1 null-mutants have a reduced ability to colonize *V. myrtillus* roots

There is increasing awareness that SSPs may play important roles during saprotrophic fungal growth, as they have been identified in saprotrophic fungi and they can be expressed by mycorrhizal fungi during asymbiotic growth (Vincent et al., 2012; Doré et al., 2015; Valette et al., 2017). However, although a limited range of growth conditions were tested, OmSSP1 did not appear to be necessary in the FLM, as the three OmSSP1-null mutants were not affected in mycelium morphology or growth rate, even when they were exposed to toxic and oxidative chemical compounds. By contrast, when they were tested for symbiotic capabilities on *V. myrtillus* plants, a significant reduction in the percentage of mycorrhization (from about 37% to about 23-24%) was measured as compared with the wild type strain, thus suggesting a specific role of OmSSP1 in the mycorrhization process. The OmSSP1 deletion did not fully prevent root mycorrhization. However, 20% of *O. maius* SSPs are induced in symbiosis, and although OmSSP1 was the most highly up-regulated, we cannot exclude a functional redundancy, as already reported for the effectors of pathogenic fungi (Selin et al., 2016). Thus, the absence of OmSSP1 could be partly compensated by other *O. maius* mycorrhiza-induced SSPs with similar function, thus lowering the impact of the OmSSP1 deletion. It will be therefore interesting to check the expression level of other symbiosis-induced OmSSPs in the OmSSP1 null-mutant strains.

## Conclusions

In conclusion, the genome of the ERM fungus *O. maius* contains several SSPs that are up-regulated in symbiosis. Decreased colonization of *V. myrtillus* roots by OmSSP1-null mutants indicates that this protein, the most highly induced in the ERM symbiosis, is a hydrophobin-like effector that participates in the molecular fungal-plant interaction occurring during mycorrhizal formation. Our data demonstrate for the first time the importance of MiSSPs in ERM, although several questions remain open on the cellular localisation of OmSSP1 and its role in symbiosis.

In ECM, hydrophobins likely play an important role in hyphal aggregation during the formation of the extraradical fungal mantle and the Hartig net (Tagu et al., 2001). However, ERM fungi do not form any extraradical hyphal aggregate on the roots of their ericaceous hosts, and individual fungal hyphae take direct contact with the hydrophilic root surface prior to cell wall penetration (Perotto et al., 2012). The different features of fungus-host plant interaction in ERM, together with the distinctive features of OmSSP1 as compared to typical Class I hydrophobins, may suggest functions specific to the ERM symbiosis.

## Acknowledgements

We thank N. Colombano for her help with some of the experiments and F. Rineau for helpful comments on the OmSSP1 biochemical features. S.C. was supported by a PhD fellowship from the Italian MIUR. The authors acknowledge financial support from local funding of the University of Turin and from the Laboratory of Excellence ARBRE (ANR-11-LABX-0002-01).

## Author Contributions

SC, SD, EMartino and SP planned and designed the research; SC, SD, EMartino, SP, AK, EMorin, H-RK, YD and CV-F performed experiments or sequencing, collected, analyzed or interpreted the data; EMartino, SP, FM contributed reagents/materials/analysis; SC, EMartino and SP wrote the manuscript; SD, CV-F and FM revised it critically. All authors read and approved the final manuscript.

